# A comprehensive analysis of vomeronasal organ transcriptome reveals variable gene expression depending on age and function in rabbits

**DOI:** 10.1101/2020.11.24.395517

**Authors:** PR Villamayor, D Robledo, C Fernández, J Gullón, L Quintela, P Sánchez-Quinteiro, P Martínez

**Affiliations:** Department of Zoology Genetics and Physical Anthropology. Faculty of Veterinary. University of Santiago de Compostela, Lugo Spain; Department of Anatomy, Animal Production and Clinical Veterinary Sciences. Faculty of Veterinary. University of Santiago de Compostela, Lugo Spain; The Roslin Institute and Royal (Dick) School of Veterinary Studies University of Edinburgh Midlothian UK; COGAL SL, Cooperativa Conejos Gallegos; Department of Animal Pathology. Faculty of Veterinary. University of Santiago de Compostela, Lugo Spain

**Keywords:** vomeronasal organ, gene expression, age-dependent changes, VNO-receptors, rabbits

## Abstract

The vomeronasal organ (VNO) is a chemosensory organ specialized in the detection of pheromones and consequently the regulation of behavioural responses mostly related to reproduction. VNO shows a broad variation on its organization, functionality and gene expression in vertebrates, and although the species analyzed to date have shown very specific features, its expression patterns have only been well-characterized in mice. Despite rabbits represent a model of chemocommunication, unfortunately no genomic studies have been performed on VNO of this species to date. The capacity of VNO to detect a great variety of different stimuli suggests a large number of genes with complex organization to support this function. Here we provide the first comprehensive gene expression analysis of the rabbit VNO through RNA-seq across different sexual maturation stages. We characterized the VNO transcriptome, updating the number of the two main vomeronasal receptor (VR) families, 129 V1R and 70 V2R. Among others, the expression of transient receptor potential channel 2 (TRPC2), a crucial cation channel generating electrical responses to sensory stimulation in vomeronasal neurons, along with the specific expression of some fomyl-peptide receptors and H2-Mv genes, both known to have specific roles in the VNO, revealed a the particular gene expression repertoire of this organ, but also its singularity in rabbits. Moreover, juvenile and adult VNO transcriptome showed consistent differences, which may indicate that these receptors are tuned to fulfill specific functions depending on maturation age. We also identified VNO-specific genes, including most VR and TRPC2, thus confirming their functional association with the VNO. Overall, these results represent the genomic baseline for future investigations which seek to understand the genetic basis of behavioural responses canalized through the VNO.

**HIGHLIGHTS:**

1. First description of the rabbit vomeronasal organ (VNO) transcriptome
2. VNO contains a unique gene repertoire depending on the species
3. High fluctuation of the VNO gene expression reveals changes dependent on age and specific functions
4. Most vomeronasal-receptors (VR) and transient receptor potential channel 2 (TRPC2) genes are VNO-specific
5. Reproduction-related genes shows a wide expression pattern

## INTRODUCTION

Many animals rely on chemical communication to regulate social and reproductive interactions between conspecifics (Wyatt, 2017). This is usually mediated by pheromones, chemical cues mainly detected by two multigenic families, vomeronasal type-1 and type-2 receptors (V1R and V2R), which are expressed in the neuroepithelium of the vomeronasal organ (VNO) (Dulac and Axel, 1995; Ryba and Tirindelli, 1997), a specialized structure located in the nasal cavity and containing vomeronasal sensory neurons (VSNs) (Barrios et al, 2014). The threshold for detecting some of these chemicals is remarkably low, near 10^−11^ M, placing VSNs among the most sensitive chemodetectors in mammals (Leinders-Zufall et al. 2000).

The VNO displays unique anatomical, physiological, biochemical and genetic features depending on the species (Salazar et al. 2016). Semiochemical perception and signal amplification and transduction from the VNO to the central circuits of the brain has been mostly studied in rodents (Isogai et al. 2012; Mohrhardt et al. 2018). However, the molecular and cellular rationale underlying vomeronasal function is not fully understood, likely due to the wide range of molecules detectable by the VNO -from proteins or peptids to steroids, histocompatibility major complex (HMC) molecules and other small metabolites (Leinders-Zufall et al. 2004; Kimoto et al. 2005; Ibarra-Soria et al. 2014a; Fu et al. 2015)–, and also to the diverse family receptor repertoire found in VSNs (V1R, V2R, formyl-peptide and HMC receptors) (Ishii et al. 2003; Revière et al. 2009; Francia et al. 2014). Additionally, the number of V1R and V2R genes varies greatly across mammalian genomes, from no functional genes (pseudogenes) in macaques to several hundreds in rodents (Young et al. 2010).

The genomic characteristics of the VNO have been studied via comparative genomics, comparing genome assemblies of different species (Grus et al. 2005), and via gene expression specific VR genes or subsets of genes by RT-PCR, *in situ* hybridization and microarrays (Zhang et al. 2010; Ishii and Mombaerts, 2011; Broad and Keverne, 2012; Kubo et al. 2016). However, only a few studies have addressed the analysis of the whole transcriptome of the VNO (Ibarra-Soria et al. 2014b; Saraiva et al. 2015; Yohe et al. 2019). Next-generation sequencing technologies can help to understand the expression profiles of the whole vomeronasal transcriptome, offering the possibility of integrating information from different transduction pathways, secondary signalling processes and regulatory interactions among VSNs.

By using RNA sequencing (RNA-seq), the full length transcripts of vomeronasal receptors were described for the first time in a mammal species by Ibarra-Soria et al. (2014b) using the mouse as a model. Saraiva et al. (2015) compared both the olfactory and vomeronasal transcriptomes of mouse to the olfactory transcriptome of zebrafish through RNA-seq, finding a very high molecular conservation between the two species. Thus far, the global VNO transcriptome has not been described in any other species excluding the recent study of the VNO transcriptome of bats (Yohe et al. 2019). Instead, effort has been invested in further RNA-seq studies in mice under different experimental conditions, disclosing variation in vomeronasal receptors expression between strains (Duyck et al. 2017), with pregnancy (Oboti et al. 2015) and under sex-separation (van der Linder et al. 2018), suggesting that VNO features may vary not only among species but also under different conditions within the same species.

Rabbits are a species pertaining to the order Lagomorpha, phylogenetically close to Rodentia. However, since pheromones are species-specific signals, each species should be considered independently according to their reproductive patterns and behavioural priorities (Brennan and Zufall, 2006). Indeed, the rabbit is the only mammalian species in which a mammary pheromone-2-methyl*-*but-2-enal (2MB2)- has been detected (Schaal et al. 2003), becoming one of the best study models of pheromonal communication in mammals (Schneider et al. 2018). Additionally, rabbits are a farm species and more recently they have also become a common pet, being the third preferred pet worldwide after dogs and cats (BCSPCA 2020); thus understanding pheromone perception by the olfactory subsystems may contribute towards the implementation of pheromone-based therapies for improving both animal production and welfare.

Certainly, some behavioural studies have been conducted to improve rabbit well-being in farms by using pheromones (Bouvier and Jacquinet, 2008), and the Rabbit Appeasing Pheromone -2MB2 analog– has been commercialized as a method to reduce stress, increase reproductive efficiency and improve animal welfare. However, these applications should be supported by comprehensive scientific studies not only based on behavioural responses to “pheromone compounds”, but also by integrating anatomical, genomic, and biochemical approaches.

Accordingly, some behavioural data regarding the rabbit mammary pheromone have been collected (Coureaud et al. 2010; Charra et al. 2013; Schneider et al. 2016), which along with a comprehensive study of the vomeronasal system at anatomical level (Villamayor et al. 2018, 2020), highlighted the relevance of chemocommunication in this species. The rabbit VNO was reported to be highly developed with many specific morphological features (Villamayor et al. 2018), but surprisingly, it has not been considered in most vomeronasal phylogenetic studies. Indeed, Grus et al. 2005 showed that the complete repertoire of rabbit vomeronasal receptors has still to be determined, and to date 160 V1R have been reported in the rabbit genome (Young et al. 2010), and 37 V2R have been shown in a phylogenetic tree of mammals by Francia et al. (2014). However, by searching in the NCBI gene database we found only 15 genes annotated as V2R in the rabbit genome assembly. Additionally, despite the well-characterized VR receptors subfamilies in mice (Tirindelli et al. 2009), no data regarding VR subfamilies and clades in rabbits have been found.

Despite our understanding of vertebrate pheromonal communication has been revolutionized by molecular-genetic approaches (Brennan and Zufall, 2006), to our knowledge there are no studies on the gene expression in the rabbit VNO using genomic approaches. A comprehensive analysis of the complete gene repertoire belonging to this organ is an essential baseline towards understanding how behaviour is wired into the central nervous system. Here, we carried out an RNA sequencing (RNA-seq) approach to: i) characterize the rabbit VNO transcriptome; ii) define the vomeronasal receptors expressed in the rabbit VNO iii) identify vomeronasal-specific genes by comparing the vomeronasal transcripts repertoire to other rabbit tissues; and iv) evaluate differences in gene expression between juveniles and adults.

Here we provide a broad outlook of the rabbit vomeronasal gene repertoire, identifying VNO-specific genes, and observing a considerably fluctuation in their overall expression profile, especially depending on age. This latter comparison has not been previously reported in any species, despite being a key biological difference since VNO activity plays a pivotal role in reproductive behaviour. Altogether results suggest a great flexibility of the vomeronasal expression in rabbits.

## MATERIAL & METHODS

### Experimental design and sampling

We employed 24 animals, separated in 12 juveniles and 12 adults (40 and 180 days, respectively) including the same number of males and females within each group. All animals were commercial hybrid-Hyplus strain PS19- and stayed on a farm (COGAL SL, Rodeiro, Spain) under the same temperature conditions (18-24°C), dark-light cycles of 12:12 hours and *ad libitum* drinking and feeding. All individuals were humanely sacrificed by an abattoir of the same company, in accordance with the current legislation, and their heads were separated from the carcasses in the slaughtering line. The whole VNOs were immediately dissected out after opening the lateral walls of the nasal cavity and removing the nasal turbinates. The hidden location of the VNOs in rabbits under the mucosal covering of the ventral part of the nasal septum, and their anatomical structure have been previously described in Villamayor et al. (2018). Previous experience was critical to obtain the organs in a short period of time (< 5 minutes after death) essential for obtaining the appropriate RNA quality (see below). Both vomeronasal organs of each animal were collected and immediately stored in Trizol and kept in ice (~4°C). Due to the double bone and cartilage envelope of the rabbit VNO, the tissue was homogenized using a mixer to guarantee the whole tissue sample was soaked by Trizol. After 20 minutes, all samples were stored at −80°C for further RNA extraction.

### Transcriptomic Analysis

RNA extraction was performed using Trizol followed by DNase treatment in accordance with the manufacturer’s instructions. RNA quantity and integrity were evaluated in a NanoDrop^®^ ND-1000 spectrophotometer (NanoDrop^®^ Technologies Inc.) and in a 2100 Bioanalyzer (Agilent Technologies), respectively. The 24 RNA samples had RNA integrity number (RIN) values > 7.6, so appropriate for library construction and sequencing. Samples were barcoded and prepared for sequencing by Novogene (Cambridge, UK) on an Illumina Nova-Seq 150 bp PE run.

The quality of the sequencing output was assessed using FastQC v.0.11.7 (https://www.bioinformatics.babraham.ac.uk/projects/fastqc/). Quality filtering and removal of residual adaptor sequences were performed on read pairs using Trimmomatic v.0.39 (Bolger et al. 2014). Specifically, residual Illumina-specific adaptors were clipped from the reads, the read trimmed if a sliding window average Phred score over five bases was < 20, and only reads where both pairs had a length longer than 50 bp post-filtering were retained. Filtered reads were aligned against the latest version of the rabbit genome (OryCun2.0; Carneiro et al. 2018) using STAR v.2.7.0e (Dovin et al. 2013) two-pass mode and the following parameters: the maximum number of mismatches for each read pair was set to 10% of trimmed read length, and minimum and maximum intron lengths were set in 20 bases and 1 Mb, respectively. Aligned reads were assigned to genes based on the latest annotation of the rabbit genome (Ensembl release 101).

Gene count data were used to calculate gene expression and estimate differential expression using the Bioconductor package DESeq2 v.1.28.1 (Love et al. 2014) in R v.3.6.2 (R Core Team 2017). Briefly, size factors were calculated for each sample using the ‘median of ratios’ method and count data were normalized to account for differences in library depth. Next, gene-wise dispersion estimates were fitted to the mean intensity using a parametric model and reduced towards the expected dispersion values. Finally, differential gene expression was evaluated using a negative binomial model that was fitted for each gene, and the significance of the coefficients was assessed using the Wald test. The Benjamini-Hochberg false discovery rate (FDR) correction for multiple tests was applied, and transcripts with FDR < 0.05 were considered differentially expressed genes (DEGs). Hierarchical clustering and principal component analyses (PCA) were performed to assess the clustering of the samples and identify potential outliers over the general gene expression background. The R packages “pheatmap”, “PCAtools” and “EnhancedVolcano” were used to plot heatmaps, principal component analysis (PCA) and volcano plots, respectively. Gene Ontology (GO) and KEGG pathway enrichment analyses were performed using the David 6.8 Bioinformatic Database (Huang et al. 2009a; Huang et al. 2009b) for all conditions.

### Vomeronasal receptors. Comparison among different rabbit transcriptome tissues

Vomeronasal receptors gene expression was compared to that of other seven rabbit tissues (hindbrain, forebrain, ovary, testis, liver, heart and kidney) in the Rabbit Expression Atlas (https://www.ebi.ac.uk/gxa/experiments/E-MTAB-6782/Supplementary%20Information; Cardoso-Moreira et al. 2019). Genes of the VNO were considered functionally relevant for the VNO when at least the expression of 22 out of 24 samples analyzed (measured as Transcripts Per Million, TPM) doubled the expression of 102 out of 104 samples of the atlas.

## RESULTS

### 1. RNA sequencing output

A total of ~843 million paired-end (PE) reads were generated, for an average of ~35 million reads for each of the 24 samples analyzed. After filtering, ~798 million reads were retained (94.7 %; > 33 million reads per sample). On average, 76.6% of the PE filtered reads were aligned to the available rabbit genome (OryCun2.0; Carneiro et al. 2018) at unique (70.0%) or multiple (6.6%) genomic positions. These reads were assigned to the 29,587 genes identified in the rabbit genome. A total of 7,690 out of the 22,675 protein coding genes in the rabbit genome did not present any significant annotation, and were blasted against the protein database Swiss-Prot (The UniProt Consortium, 2019) for updating, 2,670 additional genes being annotated.

### 2. Vomeronasal transcriptome

Given the lack of previous molecular and genetic knowledge about the rabbit VNO, we firstly characterized its global transcriptome. A total of 19,482 genes (65.8%) with at least one read were identified, 14,584 (49.3%) of them with ≥ 20 reads (Table 1); this is the threshold that we establish to consider a gene as expressed based on the distribution of the number of reads per gene (Supplementary file 1). We were especially interested in understanding the particular expression of the two main vomeronasal receptor multigene families (V1R and V2R), each of them divided into specific subsets of neurons. Most vomeronasal receptor genes showed relatively low expression, especially those pertaining to the family V1R (Table 1).

**Table 1.** Gene expression in the rabbit vomeronasal organ.

| Range of expression | Genes | VR | V1R | V2R |
| --- | --- | --- | --- | --- |
| *Whole transcriptome* | *29,587* | *199* | *129* | *70* |
| $\geq 1$ read | 19,482 | 179 | 113 | 66 |
| $\geq 20$ reads | 14,584 | 66 | 22 | 44 |
| Low expression (20-100 reads) | 3,019 | 52 | 22 | 30 |
| Moderate expression (100-1000 reads) | 8,573 | 14 | 0 | 14 |
| High expression ($\geq 1000$ reads) | 2,601 | 0 | 0 | 0 |

GO term and KEGG pathway enrichment analyses of the expressed genes (≥ 20 reads) (Supplementary file 2) revealed the over-representation of several remarkable functional categories such as PI3K-Akt signaling, an important pathway for survival and proliferation of vomeronasal sensory neurons (VSNs) (Xia et al. 2006); Transforming Growth Factor Beta Receptor, which appears to be involved in the network between the VNO and the accessory olfactory bulb (AOB) (Naik et al. 2020); and Wnt signaling, which is involved in olfactory connections and also regulates stem cell fate in the main olfactory epithelium (Zaghetto et al. 2007; Fletcher et al. 2017).

Other enriched functions were related to the maintenance of the vomeronasal sensory neurons (i.e. neuron projection, signaling pathways regulating pluripotency of stem cells and axon guidance) and to the suppression of the immune response, such as primary immunodeficiency, negative regulation of cytokine production involved in immune response or negative regulation of inflammatory response.

To refine the analyses of the rabbit VNO transcriptome constitution, we considered different gene expression ranges: i) low expression (20-100 reads), ii) moderate expression (100-1000 reads) and iii) high expression (≥ 1000 reads) (Table 1).

When assessing the functional enrichment in the three different subgroups, we found that terms related to neuron functionality were expressed mainly at low levels (i.e. presynaptic membrane, neuronal cell body membrane and glutamatergic synapse), which suggests that the ‘basic needs’ of the VSNs are fulfilled with low expression levels. Furthermore, VSNs signal transduction is mediated by G-protein coupled mechanisms, and accordingly, terms such as heterotrimeric G-protein complex or G-protein coupled receptor signaling pathway were also expressed at low levels. Our data also revealed the low expression in the VNO of ERK1 and ERK2, pertaining to a family related to one of the most prominent intracellular cascades, which links extracellular G protein coupled receptor stimulation with the activation of effector proteins. ERK1/2, also known as pMAPK, is the phosphorylated form of the mitogen-activated protein kinase (MAPK). Both MAPK and ERK1/2 have been involved in the activation of VSNs (Dudley et al. 2001). Other pathways such as chemoattractant activity or steroid binding were also expressed at low level and might be related to the VNO capacity to response to external stimulus and to mediate reproductive behaviour, respectively.

Moderate and high expression genes were instead mainly enriched in reproduction pathways either directly (i.e. GnRH, estrogen, oxytocin, prolactin signaling pathways and progesterone-mediated oocyte maturation) or indirectly (i.e. circadian rhythm, importantly involved in seasonal reproduction of animals, and p53 signaling pathway, whose loss leads to a significant decrease of fertility). Other pathways found at those expression ranges are related to VSNs involving growth and maintenance (Neurotrophin signaling pathway), differentiation and regeneration (Notch signaling pathway) and migration and axon guidance.

### 3. Vomeronasal receptors

We identified a total of 199 VR genes in the rabbit genome, which is in the upper range of the broad VR gene repertoire of mammals, from > 250 intact (likely functional) VR in mice to ~40 intact VR in cows and ~8 in dogs. More specifically, we updated the number of VR for each of the two main vomeronasal families, finding 129 V1R and 70 V2R genes in the rabbit genome (Supplementary file 3). The latter figure is rather surprising because we identified up to 55 new V2R, a family completely degenerated in most mammals. Additionally, following the same criteria as Dinka et al. 2016, we classified the total VR genes repertoire into 177 intact VR (encoding > 300 amino acids (aas)) and 22 partial VR (> 100 aas and < 300 aas).

Despite the vast majority of vomeronasal receptors, especially V1R, showed relatively low expression (Table 1), a large dynamic range of expression was observed in the VRs identified (Figure 1). The highest expressed gene was VN2R1 (ENSOCUG00000002314) with 308.20 normalised reads, while 14 V2R genes showed moderate expression levels (100-1000 reads). However, most V2R were lowly expressed (20-100 reads), and the majority of V1R genes showed less than 20 reads on average (Supplementary file 3). The mean expression for V1R was 11.48 whereas for V2R was 54.47. This is not surprising because most V1R are thought to be expressed only in one or two specific subsets of neurons, being tuned up by a cognate ligand, while at least some V2Rs can be co-expressed in the same neurons (Martini et al. 2001).

**Figure 1.**
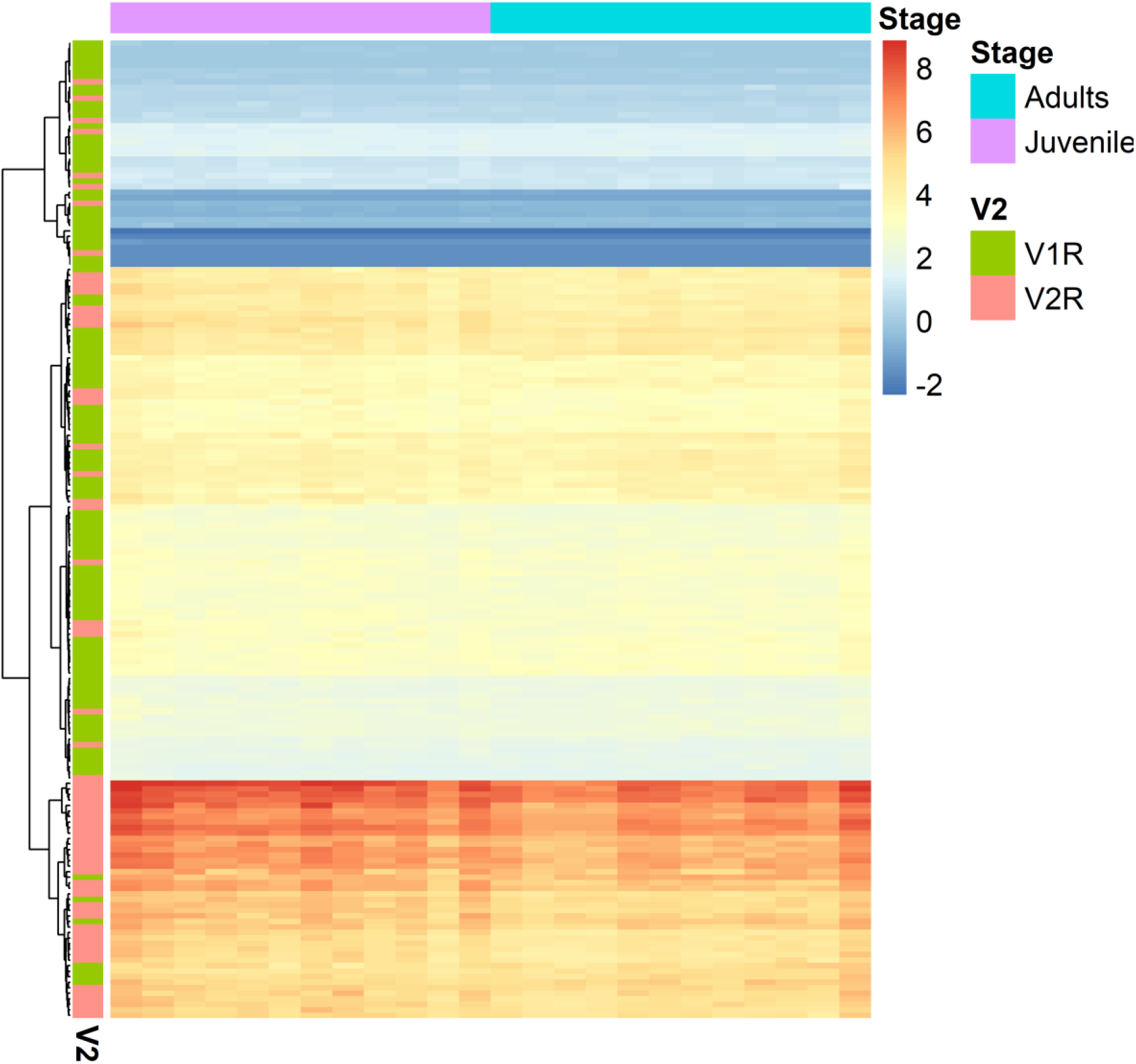
Expression of vomeronasal receptor genes in the rabbit vomeronasal organ.

Functional enrichment analysis of the VR (Supplementary file 3) revealed only two enriched GO terms in the molecular function category: pheromone receptor activity and G-protein coupled receptor activity, both linked to pheromone detection which is the main role of VSNs. Neither biological process nor KEGG pathway significant enrichment was found, but three important protein domains were detected: Vomeronasal type 1 receptor family signature, Vomeronasal type 2 receptor family signature and Metabotropic glutamate GPCR signature.

A third class of vomeronasal receptors is the formyl peptide receptor (FPR) family, previously reported in mice (Liberles et al. 2009; Rivière et al. 2009). In rabbits, two genes from this family were found in the VNO (FPR1 and FPR2) with low expression and no differences between ages. Another multigene family, named H2-Mv and belonging to the MHC, was previously reported in neurons of the VNO basal layer in mice (Ishii et al. 2003). We blasted these genes against the rabbit genome and, from the 9 H2-Mv identified in mice, we detected 6 in rabbit. All of them were also found in our VNO transcriptome (Supplementary file 6, HMC), being mostly highly expressed and showing no differences between ages.

### 4. Vomeronasal-specific gene expression

To assess whether the genes of the rabbit vomeronasal organ are VNO-specific, we compared their expression in this organ to that in hindbrain, forebrain, ovary, testis, liver, heart and kidney using publicly available RNA-Seq data (Cardoso-Moreira et al. 2019) (Supplementary file 7). We found 429 genes that are consistently VNO-specific. Among them, 80 vomeronasal receptors (45 V1R and 35 V2R) were considered VNO-specific, whereas 71 VR (47 V1R and 24 V2R) were not found in the rabbit expression atlas, and therefore their VNO-specificity could not be assessed. Additionally, some VR VNO-specific, especially V1R, showed high expression in the adult testis (Supplementary figures 1 and 2). We then analyzed the expression of the remaining 48 VR (37 V1R and 11 V2R) supposed to be ‘non VNO-specific’, and interestingly, their overall expression was higher in the VNO than in the other analyzed tissues (Supplementary figures 3 and 4). However, in some samples their expression was 0, which hinder the performance of a thorough statistical analysis. Additionally, some VR belonging to the ‘non VNO-specific’ group, were slightly expressed in other tissues, especially in testis, ovary and brain. No expression of VR genes was found in heart, liver, and kidney (Supplementary figures 1-4).

The transient receptor potential channel 2 gene (TRPC2) and four out of six genes from the H2-Mv family were also VNO-specific. Additionally, we added to this category six other genes of the MHC, either from class I or class II, as well as several immunoglobulins and other immunity-related genes such as lipocalins. More in detail, the major urinary protein IV (MUP4) and other two lipocalin (ENSOCUG00000026763, ENSOCUG00000028189; both of them coded as LCN1) were among the most expressed genes in the VNO (>164,191 reads) and considered VNO-specific, thus suggesting the importance of this proteins in the VNO function.

Other genes were found to be expressed substantially higher in the VNO than in other tissues. For example, Golgi associated olfactory signaling regulator (GFY), which is expressed in both the VNO and the main olfactory epithelium (MOE) throughout development in mice (Kaneko-Goto et al. 2013); or receptor transporter protein 2 (RTP2), which enhances the response of olfactory receptors (OR) and previously reported to be specific of olfactory neurons (Saito et al. 2004). Furthermore, two prolactin and one androgen related-genes were also VNO-specific, but no other reproductive-related genes were found, which is consistent with the well-known expression of these genes in endocrine tissues and neural circuits (Inoue et al. 2019).

Looking at the molecular function, the terms ‘pheromone receptor activity’, ‘peptide antigen binding’, ‘G-protein coupled receptor activity’ and ‘transmembrane signaling receptor activity’ were enriched in vomeronasal-specific genes, which supports pheromone-detection as its principal role. Additionally, some biological process and cellular component terms were related to the immune system (i.e. immune response, antigen processing and presentation and some MHC related-terms), as expected considering the wide range of immune-related genes found in the list (Supplementary file 7). This is consistent with previous studies where genes from immune response and chemosensory receptor classes were found as the most represented genes in the VNO (Duyck et al. 2017).

### 5. Differential expression of the vomeronasal organ between juveniles and adults

The vomeronasal organ transcriptome showed sharp differences between juvenile and adult rabbits, with both groups showing somewhat dispersed clustering in two well differentiated groups in the PCA (Figure 2) and volcano plot (Figure 3). A total of 3,061 DEG were detected between juveniles and adults, 1,376 up-regulated (higher expression in adults) and 1,685 down-regulated (higher expression in juveniles) (Supplementary file 4). Some genes of the VR repertoire were down-regulated, including six and one genes of the V2R and V1R families, respectively. Additionally, several genes related to vomeronasal function were either up- or down-regulated, such as SEMA3F, KIRREL2, ANO10 and ROBO2. Interestingly, TRPC2, essential in VNO signal transduction, was up-regulated, whereas TRPC1 was down-regulated. Other groups such as arginase (ARG1, ARG2) or Notch-related genes (Delta-like 4 signaling gene (Dll4), Delta like non-canonical Notch ligand 1 (DLK1)) also showed differences between ages. All in all, a set of 32 genes would enable the classification of all individuals in our experiment with full confidence (Figure 4).

**Figure 2.**
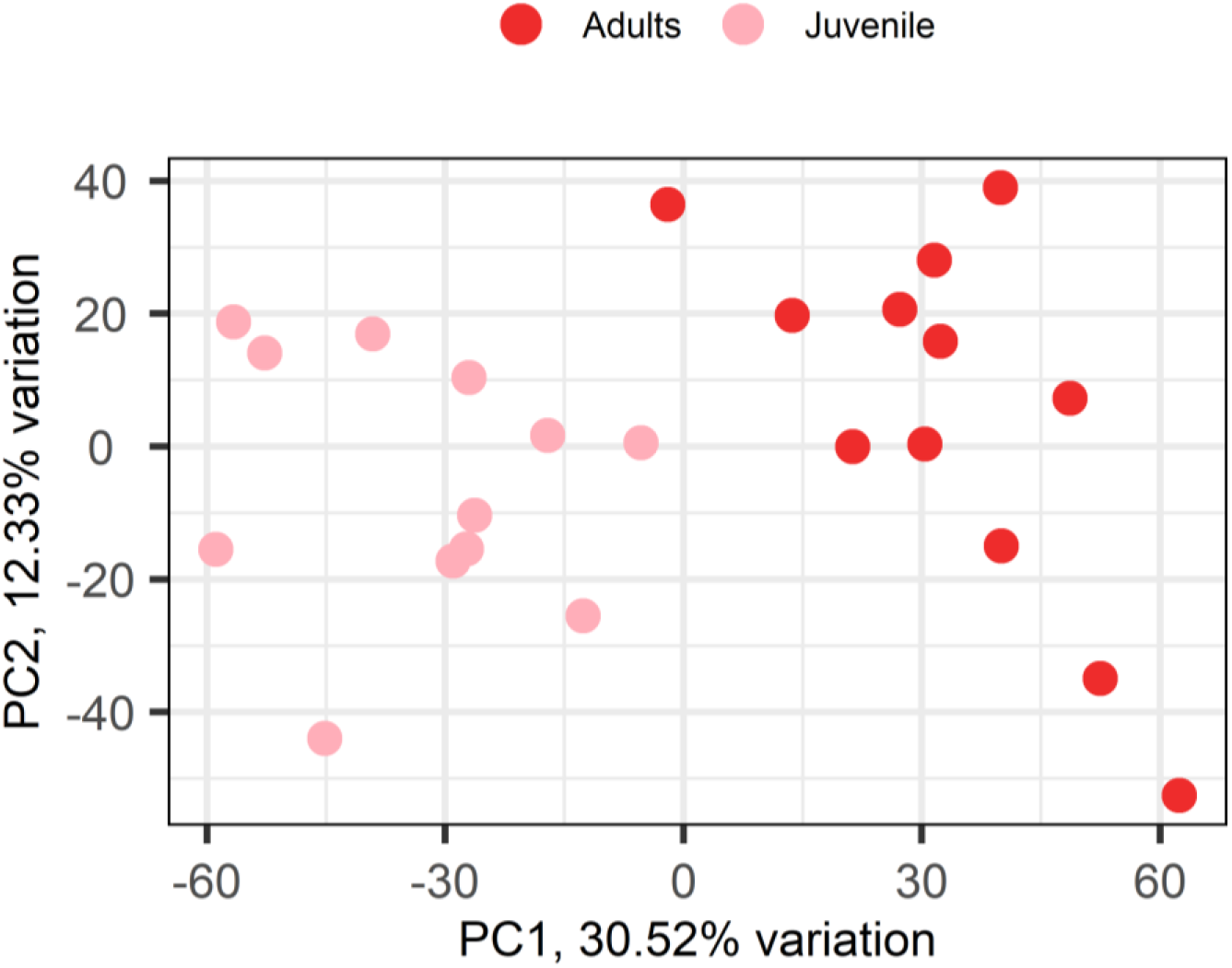
Principal component analysis of the vomeronasal organ transcriptome.

**Figure 3.**
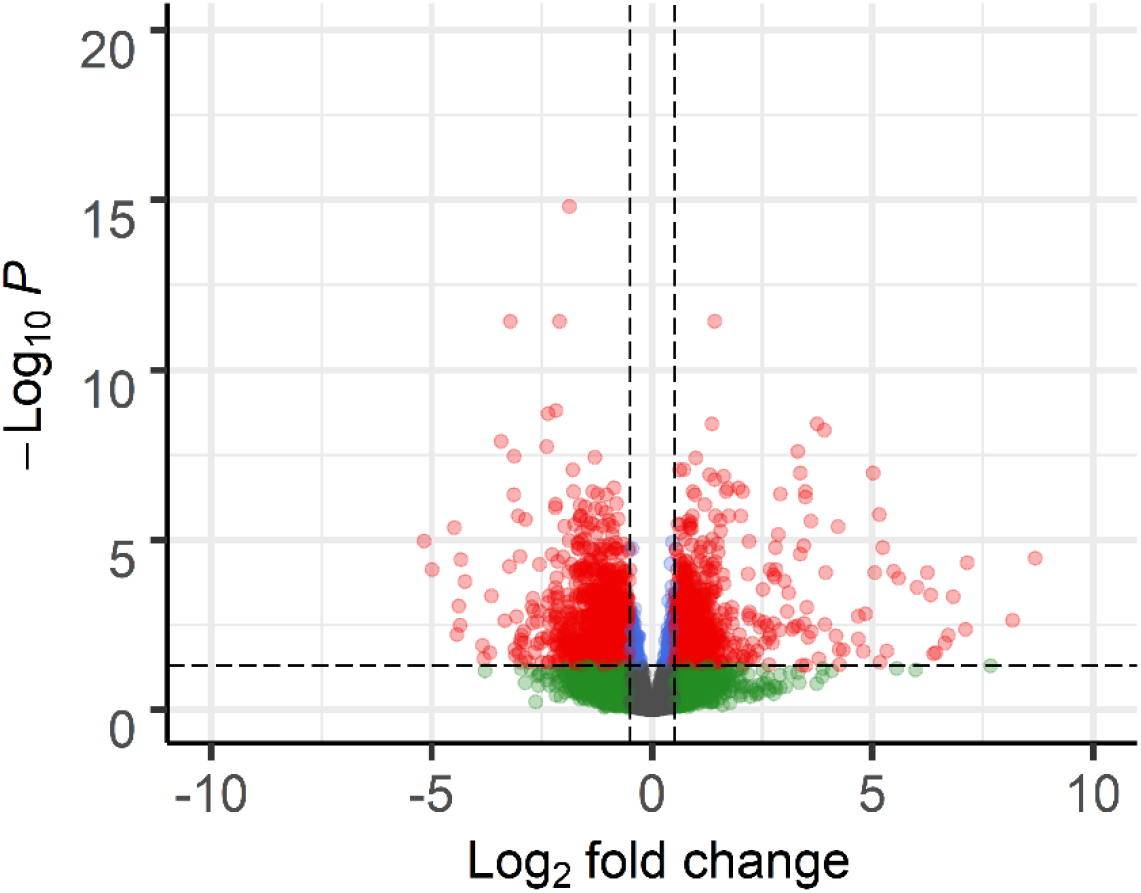
Volcano plot of the vomeronasal organ transcriptome.

**Figure 4.**
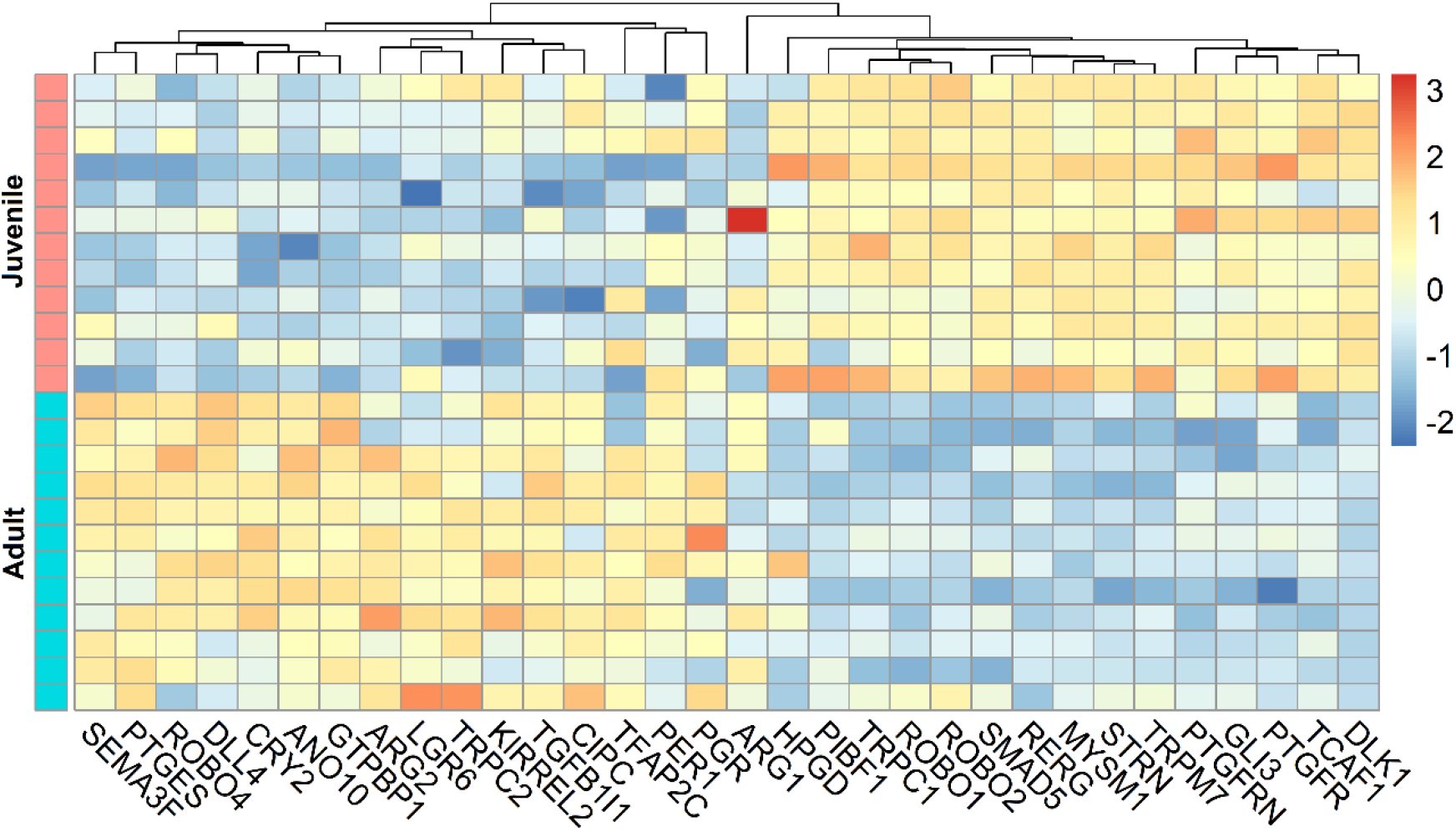
Differential expression between the vomeronasal organ of juvenile and adult rabbits.

GO term and KEGG pathway enrichment analyses were performed for both lists of up-regulated (more expressed in adults) and down-regulated (more expressed in juveniles) genes separately (Table 2; Supplementary file 5). In juveniles, terms such as dendritic morphology and axon guidance, putatively related to establishing olfactory connections, were enriched, which is further supported by the over-expression of Wnt signaling pathway. Instead, adults showed enriched terms such as receptor activity, neuronal cell body and myelination in peripheral nervous system, which fits the functional role of the adult vomeronasal organ.

**Table 2.** List of some highlighted up- and down-regulated GO terms and KEGG pathways when comparing juvenile and adult vomeronasal organ.

| CATEGORIE | TERM | NUMBER OF<br>GENES | P-VALUE<br>Benjamini |
| --- | --- | --- | --- |
| **UP-REGULATED** |  |  |  |
| Biological Process<br>(BP) | Circadian regulation<br>of gene expression | 9 | 1,00E+00 |
| BP | Entrainment of<br>circadian clock by<br>photoperiod | 5 | 1,00E+00 |
| BP | Negative regulation of<br>ERK1 and ERK2<br>cascade | 11 | 1,00E+00 |
| BP | Notch receptor<br>processing | 3 | 1,00E+00 |
| BP | Negative regulation of<br>Notch signaling<br>pathway | 7 | 9,90E-01 |
| BP | Negative regulation of<br>neuron projection<br>development | 7 | 9,90E-01 |
| BP | Myelination in<br>peripheral nervous<br>system | 4 | 1,00E+00 |
| Cellular Component<br>(CC) | Neuronal cell body | 17 | 7,50E-01 |
| CC | Myelin sheath | 33 | 2,90E-03 |
| Molecular Function<br>(MF) | Receptor activity | 11 | 9,90E-01 |
| KEEG Pathway | Estrogen signaling<br>pathway | 16 | 3,80E-01 |
| **DOWN-REGULATED** |  |  |  |
| BP | Circadian regulation<br>of gene expression | 12 | 9,80E-01 |
| BP | Regulation of<br>circadian rhythm | 8 | 1,00E+00 |
| BP | ERK1 and ERK2<br>cascade | 6 | 9,90E-01 |
| BP | Negative regulation of<br>canonical Wnt<br>signaling pathway | 18 | 9,90E-01 |
| BP | Dendrite<br>morphogenesis | 9 | 9,90E-01 |
| BP | Stem cell population<br>maintenance | 11 | 9,90E-01 |
| KEEG Pathway | Wnt signaling<br>pathway | 21 | 2,00E-01 |
| KEEG Pathway | PI3K-Akt signaling<br>pathway | 48 | 3,40E-02 |
| KEEG Pathway | MAPK signaling<br>pathway | 39 | 3,30E-02 |
| KEEG Pathway | p53 signaling pathway | 19 | 5,70E-03 |
| KEEG Pathway | Signaling pathways<br>regulating<br>pluripotency of stem<br>cells | 21 | 1,90E-01 |
| KEEG Pathway | Axon guidance | 20 | 2,00E-01 |
| KEEG Pathway | Progesterone-mediated oocyte<br>maturation | 16 | 1,60E-01 |
| KEEG Pathway | GnRH signaling<br>pathway | 17 | 7,20E-02 |
| KEEG Pathway | Prolactin signaling<br>pathway | 11 | 4,00E-01 |

### 6. Wide expression of reproduction-related genes

Regarding reproduction, several terms (i.e. estrogens, gonadotropin releasing hormone (GnRH), circadian clock) and genes (i.e. prolactin (PRLR, PIP) or progesterone (PIBF1, PGR)) were either found in juveniles or adults suggesting a relationship between the VNO and this biological process from the first life-stages. Accordingly, the terms progesterone, GnRH and prolactin were found in the juveniles list of enriched KEGG pathways (Table 2; Supplementary file 5). Eight progesterone-related genes (1 up- and 1 down-regulated) and three prolactin-related genes were detected in the rabbit VNO transcriptome (see Supplementary file 6 for gene-groups). The case of GnRH is interesting because only two receptors (GNRH1 and GNRHR) with very low expression-between 1-10 reads, which is below the established threshold (see M&M)- were found. However, other genes which control GnRH production were moderately expressed in the rabbit VNO such as Gli3, down-regulated and related to GnRH-1 migration to the central nervous system (Taroc et al. 2020); and Dmxl2 and Lgr4, moderately expressed in our samples and whose deficiency results in decreased fertility or delayed puberty due to fewer GnRH neurons (Tata et al. 2014; Mancini et al. 2020).

Additionally, the term estrogens was found in the adult list of GO and KEGG pathways (Table 2; Supplementary file 5), and from the nine estrogen-related genes detected, only two were DEG (RERG and STRN), being down-regulated. Moreover, 15 genes related to androgens (1 up- and 1 down-regulated) were moderately expressed on average (Supplementary file 6), whereas oxytocin receptor (Oxtr), luteinizing hormone/choriogonadotropin receptor (Lhcgr) and follicle-stimulating hormone receptor (Fshr) showed lower expression or were not detected. We also found 34 genes related to spermatogenesis, 5 and 11 up- and down-regulated, respectively, but their function in the VNO remains unknown. Finally, other pathway related to reproduction and found in both lists of juveniles and adults was the circadian clock, which supports the importance of the photoperiod in this species. We detected 11 genes involved in circadian rhythm, four of them up-regulated (Table 2; Supplementary file 5 and 6).

As expected, we found relevant differences between juvenile and adult VNO. To our knowledge this is the first time in which a comparative transcriptomic analysis between these two groups has been performed in the VNO. Our data demonstrate that the rabbit VNO is fully active 40 days after birth (just post-weaning), expressing several reproduction-related genes. However, the VNO increases its activity in mature animals (6 months old) supported by up-regulated terms such as “receptor activity” as well as the over-expression of TRPC2 in adults.

## DISCUSSION

This study provides a comprehensive gene expression analysis of the rabbit vomeronasal organ through RNA-seq, characterizing for the first time the rabbit VNO transcriptome. Remarkably, the two main VR families (V1R and V2R) showed very low expression, especially V1R, thought to express a single sensory receptor in mice (Belluscio et al. 1999; Rodriguez et al. 1999), thus supporting a ‘one cell-one receptor’ model in the VNO (Luo and Katz, 2004). We also identified VNO-specific genes, when their expression doubles the expression in 22 out of the 24 analyzed tissues. Moreover, juvenile and adult VNO transcriptome comparison was described here for the first time, revealing considerable fluctuation in the vomeronasal repertoire depending on age.

### 1) New strategy to approach vomeronasal gene expression

The vomeronasal organ acts as an interface between the immune and nervous systems and carries out many functions such as pheromone perception and signal transduction, receptor activity and neuron maintenance, allowing the regulation of a wide range of behaviours, including individual recognition, reproduction, and aggression (Brennan 2004; Chamero et al. 2012; Francia et al. 2014; Brignall and Cloutier, 2015). This is mediated by highly qualified sensory receptors expressed in the neuroepithelium of the VNO, whose high specificity results in very low expression. Due to the various functions of the VNO, it is reasonable that its transcriptome shows wide intrinsic variability and varies over time depending on the function at each life-stage and social interaction. Accordingly, a new approach to unveil the whole vomeronasal transcriptome has been addressed here, considering low (20-100 read), moderate (100-1000 reads) and high (> 1000 reads) expressed genes.

### 2) The unique vomeronasal gene repertoire

#### a) Vomeronasal-type receptors

The rapid evolution of the two main vomeronasal receptor superfamilies -V1R and V2R- in some genome lineages place them among those gene families showing the broadest gene number variation (Shi and Zhang, 2007; Nei et al. 2008). For instance, snakes and lizards exhibit a large number of V2R genes, but retain an extremely limited number of V1R (Brykczynska et al. 2013). Instead, mammals with a highly developed VNO tend to expand V1R repertoire, as it occurs with platypus (283 genes), mouse (239), rat (109) and rabbit (here updated to 129). Intact 121 and 37 V2R (here updated to 70) were found in mouse and rabbit respectively, whereas no functional V2R genes have been reported in species such as Old World monkeys or humans (Francia et al. 2014). The loss of these receptors is confirmed by the high number of pseudogenes and a vestigial and presumably non-functional VNO (Rodriguez and Mombaerts, 2002), even though this latter is still a matter of controversy (McGann, 2017; D’Aniello et al. 2017; Salazar et al. 2019).

We therefore emphasize that the analysis of the vomeronasal gene repertoire at both genomic and transcriptomic levels should be considered in each species and each condition independently. In rabbits, in addition to the particular number of V1R and V2R in the gene repertoire, seven VR were differentially expressed between juveniles and adults (down-regulated), suggesting an active organ from the first month of life. To our knowledge, this is the first time in which a transcriptomic analysis between juveniles and adults has been performed.

On the other hand, a comparison of the rabbit VR gene expression among different tissues has revealed 80 VR VNO-specific and 48 VR ‘non VNO-specific’. This latter was not considered VNO-specific due to the criteria employed in the analysis, more than biological reasons (see results), and in both cases VR genes showed overall higher expression in the VNO than in other tissues. This is consistent with previous studies in mice, where the vast majority of VR genes were only expressed in the VNO (Zhang et al. 2010).

Additionally, some VR VNO-specific, especially V1R, have also been expressed in adult testis, which is not surprising since V1R genes were previously found in mouse testis (Tatsura et al. 2001). However, we can argue that in mice VR expression was found in testis developing germ cells, whereas in rabbits no expression was noticed from the embryonic day 12 to the postnatal day 80, thus being concentrated in mature animals (186-548 days). In swine, V1R genes were highly expressed in the VNO and in testis, whereas their expression in other tissues was very low (Dinka et al. 2016). The function of V1Rs in testis remains still unknown, but other chemical receptors such as olfactory receptors (ORs) were also expressed in this gonad in mice, rats and humans (Spehr et al. 2003; Makeyeva et al. 2020), thus suggesting another functional roles other than stimulating sensory systems.

Finally, in rabbits, some VR belonging to the ‘non VNO-specific’ group, were also found lowly expressed in both gonads and brain. Expression in brain and bulb was previously detected in mice (Zhang et al. 2010). No other tissues in any species have shown additional VR expression apart from the V1R expressed in the MOE of goats, mice, lemur (also showed VN2R2 expression) and humans (Rodriguez and Mombaerts, 2002; Wakabayashi, 2002; Hohenbrink et al, 2014). In rabbits, MOE transcriptome has not yet been performed, and further studies are needed to obtain a map of its chemosensory receptors expression, verifying whether VR play a role in the main olfactory network.

#### b) Formyl peptide receptors

Formyl peptide receptor (FPR) family was firstly detected in immune cells (Le et al. 2002; Silva et al. 2020) and it has also been reported as a third family of VNRs in mice (Liberles et al. 2009; Rivière et al. 2009) with up to 12 FPR genes, including 6 pseudogenes, exclusively observed in mice vomeronasal tissue extracts -the families Fpr-rs1, Fprrs3, Fpr-rs4, Fpr-rs6 and Fpr-rs7-. However, the only two FPR genes found in the rabbit vomeronasal organ transcriptome were FPR1 and FPR2 -also called Fpr-rs2– and were not transcribed in mouse vomeronasal neurons (Rivière et al. 2009), but in other cell types and tissues more closely related to immunity (Migeotte et al. 2006). Despite further studies in rabbits are needed to determine the presence of FPR1 and FPR2 in immune cell-types as well as to rule out its presence in VSNs, it is possible that the expansion of the FPR gene family encompassing an olfactory function is rodent-specific, which is also supported by previous studies in primates which only found FPR1 and FPR2 in the genome (Yang and Shi, 2010).

#### c) H2-Mv receptors (major histocompatibility complex)

In mice, a subset of VSNs, whose bodies belong to the basal layer of the VNO epithelium, coexpress Vmn2r G-protein-coupled receptor genes with H2-Mv genes (Ishii et al. 2003), an additional multigene family which represents non-classical class I genes of the major histocompatibility complex. Despite the physiological roles of the H2-Mv gene family remain mysterious, Zufall et al. (2014) showed that these genes are required for ultrasensitive chemodetection by a subset of VSNs. In rabbits, we found six H2-Mv in the VNO transcriptome, which correlates with nine of mice. However, further analyses are required to determine whether they are differentially expressed in subsets of neurons as reported in mice.

#### d) Transient receptor potential channel. Importance of TRPC2

Pheromone signals enter the vomeronasal organ and activate essential components of the signal transduction machinery. Among them, the transient receptor potential channel 2 gene (TRPC2) is crucial for the generation of electrical responses in VSNs to sensory stimulation (Zufall, 2005). We found that its expression is higher in adult than in juvenile rabbits suggesting a greater pheromone-signalling involving this cation channel in mature animals. Additionally, TRPC2 was considered VNO-specific in rabbits, which can be compared with its ‘exclusive expression’ in the VNO of rats (Löf et al 2011). Conversely, in mice TRPC2 was not only found in the VNO but also in testis, sperm, and both in the dorsal root ganglion and in the brain (Löf et al 2011). More recently, there have also been proved its presence in olfactory sensory neurons (OSNs) within the MOE in mice and lemur (Omura and Mombaerts, 2014; Hohenbrink et al, 2014). Since the MOE was not considered among the tissues that we analyzed for the VNO-specific genes, further studies regarding TRPC2 expression in the rabbit MOE would aid to elucidate whether this gene is also involved in olfactory signaling through the main olfactory pathway in rabbits.

A second transient receptor potential channel, TRPM4, has been recently involved in vomeronasal circuitry in mice, being present in VSNs (Eckstein et al. 2020); however, we did not found this receptor in the rabbit genome, and more in-depth studies would help determine its role in rabbit vomeronasal function. In contrast, we have found three TRP - TRPM7, TRPM8 and TRPC1- not previously reported as expressed in the VNO, over-expressed in juvenile rabbits, and some other TRP were lowly or moderately expressed but did not show differences between ages. Taken together, different members of the TRP family may be implicated in pheromone-signalling in both juvenile and adults.

#### e) Lipocalin family. MUP-4 in the rabbit nasal mucosa

Lipocalin (LCN) family is involved in immune response and pheromone transport (Dominguez-Pérez et al. 2019). It contains evolutionary conserved genes and almost all of them are present in all animal kingdoms (Charkoftaki et al. 2019). More in detail, the major urinary proteins (MUPs), a specific cluster within the LCN family, has been comprehensively characterized and annotated in mice by Logan et al. 2008, and the same authors described orthologous in rat, horse and grey mouse lemur, therefore considering remarkable lineage specificity, which may depend on species-specific function in mammals.

MUPs act as genetically encoded pheromones, and are involved in vomeronasal stimuli and communication of information in urine-derived scent marks (Papes et al. 2010; Kaur et al. 2014). They are mostly found in urine, liver, and exocrine glands, but interestingly, MUP-4 is an isoform known to be expressed in the vomeronasal mucosa and it is highly specific for the male mouse pheromone 2-sec-butyl-4,5-dihydrothiazole (SBT), which promotes aggression amongst males while inducing synchronized estrus in females (Sharrow et al. 2002; Perez-Miller et al. 2010).

MUPs expression has been detected in urine and liver under different conditions in mice (Stopková et al 2007; Nelson et al 2013, 2015), and more recently Unsworth et al. (2017) reported no detectable levels of MUPs in mouse lemur urine. Additionally, the expression of lipocalins on the vomeronasal organ has been longely reported (Miyawaki et al. 1994). In fact, Broad and Keverne (2012) found MUP 1-3 and 5 in the glandular region of the mice VNO, thus suggesting a role of nasal MUPs in binding pheromones and likely presenting them to their receptors. They may also provide an additional means of selectively modulating the activation of vomeronasal receptors.

Here we provide evidence that several lipocalin, including MUP-4, belong to the most expressed genes in the rabbit VNO transcriptome and they are also VNO-specific. This is the first time in which expression of a MUP is found in a non-rodent species, and since SBT nicely binds to MUP-4 in mice (Perez-Miller et al. 2010), the next step would be determining whether this compound has a pheromone-activity in rabbits. Additionally, further studies of the vomeronasal receptors involved in such signal detection would provide useful support to future functional and evolutionary studies on chemocommunication.

### 3) Insights into vomeronasal connectivity

Sensory systems are responsible of encoding information from the environment to the central nervous system. Specifically, the VNO is implicated in transforming chemical cues into electrical signals, in which the ion channels anoctamin 1 and 2 (ANO1 and ANO2) play a fundamental role (Dauner et al. 2012). These genes also contribute to vomeronasal signal amplification (Münch et al. 2018), and we found both of them expressed in the rabbit VNO. Additionally, ANO10 was up-regulated in our samples, an observation not previously reported in any species.

The functional maturation and connectivity of basal VSNs in mice is driven by Smad4 through the canonical BMP/TGFβ signaling pathway (Naik et al. 2020). We found this gene as the highest expressed one (> 773 reads) among the 8 belonging to Smad family in the rabbit VNO transcriptome. Additionally, 21 BMP and 22 TGFβ genes were expressed in the rabbit VNO, seven and three of them differentially expressed between juveniles and adults, respectively. It is remarkable the expression of BMP4, among the highest expressed genes within the BMP repertoire, which has a role in neurogenic fate for developing VSNs (Forni et al. 2013). BMP has also proved to be critical in the generation of OSNs in the MOE and development of the olfactory nerve, and its balance with Notch signaling is critical in neurogenesis regulation (Maier et al. 2011).

Despite OSNs plasticity has been more broadly studied, the VSNs of the VNO also regenerate throughout life (Brann and Firestein, 2014). In rabbits, the constant regeneration and differentiation of VSNs is reflected by stem cell and Notch-related enriched terms. In fact, we found 11 Notch genes differentially expressed, with Delta-like 4 signaling gene (Dll4) and delta like non-canonical Notch ligand 1 (DLK1) being up- and down-regulated, respectively. Dll4 is known to be expressed in the sensory epithelium of the mice VNO across development among many other tissues, suggesting multiple developmental roles (Benedito and Duarte, 2005). Additionally, Notch1 has been previously involved in differentiation and regeneration of mice VSNs (Wakabayashi and Ichikawa, 2006), but this gene was not found in the rabbit genome. Instead, Notch2 and Notch3 were moderately and highly expressed in the rabbit VNO, respectively, and they did not show differences between juveniles and adults.

The VSNs contribute to the accessory olfactory network, integrating the signals coming from the outside world and therefore contributing to the response formation in high central circuits. Robo2 is a critical gene for targeting basal VSNs axons to the posterior accessory olfactory bulb (AOB) (Prince et al. 2009), and it was found down-regulated in rabbits. Other genes from this family, Robo1 and Robo4, were down- and up-regulated respectively. Additionally, Semaphorin3F and Kirrel2, related to vomeronasal axon fasciculation and synaptogenesis to AOB, respectively (Cloutier et al. 2004, Brignall et al. 2018), were both up-regulated in the rabbit transcriptome. Within the Kirrel family of transmembrane proteins, Kirrel2 and Kirrel3 were found moderately expressed in the rabbit VNO, and both are essential for glomeruli formation in the AOB. Remarkably, they have been expressed in non-overlapping subpopulations of VSNs in mice, and their expression is regulated by the VNO activity (Prince et al. 2013; Brignall et al. 2018).

### 4) The reproductive role of the vomeronasal organ

The vomeronasal organ is involved in species-specific behaviour and plays an important role in the reproductive function (i.e sexual attraction or maternal aggression) (Martín-Sánchez et al. 2015). The transcriptomic analysis performed in this study showed several enriched GO terms and KEGG pathways as well as hormone receptor genes closely related to the own reproductive physiology of the organ.

Progesterone receptors membrane component 1 and 2 (Pgrmc1 and Pgrmc2) showed high levels of expression whereas PGR was lowly expressed. This is consistent with previous studies of the VNO in pregnant female mice (Oboti et al. 2015). We also found that PGR was up-regulated in rabbits, suggesting a broader function of this receptor in mature animals. Notwithstanding, since hormones such as progesterone may modulate sensory inputs to the VNO depending on the state of ovulation in mice (Dey et al. 2015), a comprehensive study of the activation/silencing of VSNs under progesterone exposure across rabbits ovulation cycle would help understand the mechanisms underlying state-specific behaviour in this species.

Interestingly, estrogen receptor 1 (Esr1) was poorly expressed in our samples, which contrast with the prominent expression found in the VNO of female mice (Oboti et al. 2015). Administration of estrogen to these animals is sufficient to stimulate vomeronasal progenitor cell proliferation in the VNO epithelium, likely through activation of Esr1. Further studies considering only rabbit females may help elucidate the importance of this receptor in estrogen-detection in this species.

More specifically, Gli3, a gene that control GnRH production, was moderately expressed in the rabbit VNO. This is consistent with its expression in proliferative progenitors in the apical portions of the developing mice VNO and could be related to asserting control of pubertal onset and fertility (Taroc et al. 2020). Additionally, we found moderately expressed genes such as Dmxl2 and Lgr4, which might be related not only to fertility onset but also to its maintenance throughout reproductive life (Tata et al. 2014; Mancini et al. 2020). Dmxl2 has been previously involved in mice olfactory mucosa and Dmxl2-knockouts have shown deficiencies in olfactory signal transmission (Tata et al. 2014). However, this is the first time in which its expression in the VNO is shown. Interestingly, Dmxl2 exerts its principal function in the testes at the onset of puberty and was previously found highly expressed in testis (spermatogonia and spermatocytes), therefore playing a dual role in olfactory information and first wave of spermatogenesis (Gobé et al. 2019).

Lgr4 is strongly expressed in the VNO during development in mice, and its deficiency results in impaired GnRH neuron development and delayed puberty (Mancini et al. 2020). GnRH neurons migrate along axons of cells that reside within the VNO to the forebrain, and therefore the expression of Lgr4 in the VNO may be related to the GnRH repertoire formation. Accordingly, we found this gene enriched in rabbit juveniles. Moreover, its activation enhances the canonical Wnt signaling pathway, an enriched term in our analysis in juveniles. Another gene of this family, Lgr5, was found moderately expressed in the rabbit VNO both in adults and juveniles. Lgr5 has been previously found in MOE and taste stem cells (Qin et al. 2018), but to our knowledge this is the first time in which its expression in the VNO is reported.

All in all, we have provided here a comprehensive analysis of the rabbit VNO transcriptome, considering different conditions -DE between juveniles and adults and VNO-specific genes–. Fluctuation of VR expression levels over time may indicate that these receptors are tuned to fulfill specific functions depending on the age. We have also offered a wide panorama of the vomeronasal gene repertoire in this species -VR, FPR, H2-Mv, TRPC2 among others-. Taking into account the great variability of chemical cues that animals are exposed to, as well as the flexibility observed in the vomeronasal system, there may be species-specific additional vomeronasal families essential for species survival but whose genome sequences are yet to be discovered. Finally, our results represent the baseline for future investigations which seek to understand the genetic basis of behavioural responses.

