## Supplementary figures and images for "A comprehensive analysis of vomeronasal organ transcriptome reveals variable gene expression depending on age and function in rabbits"

### Supplementary figure 2

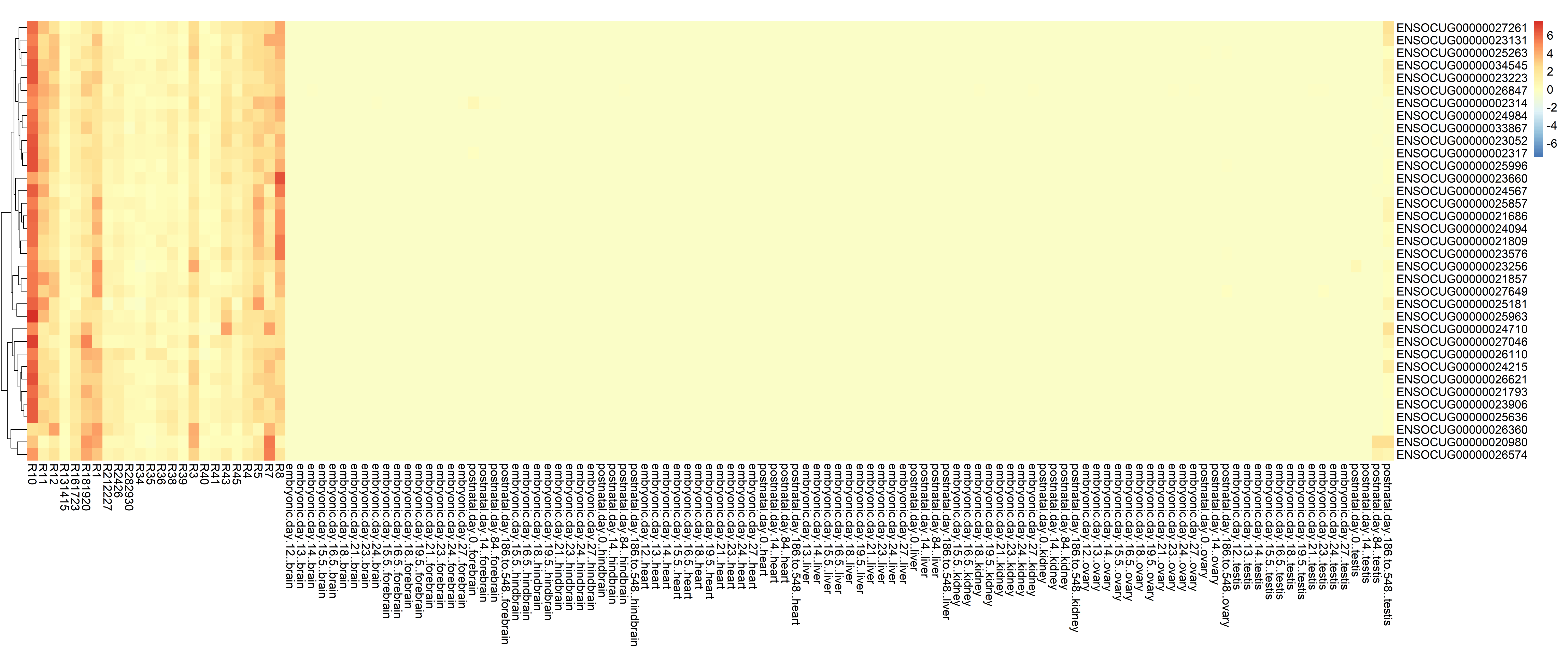

### Supplementary figure 3

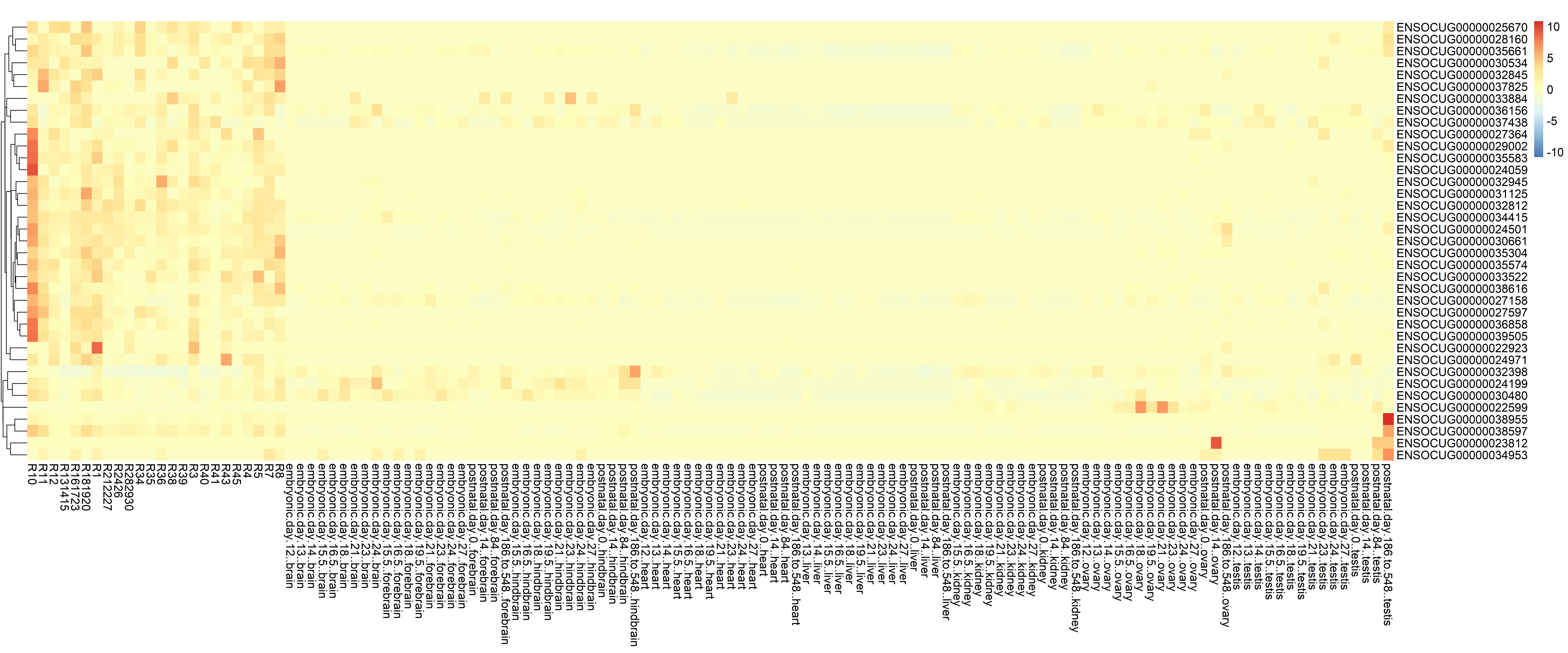

### Supplementary figure 4

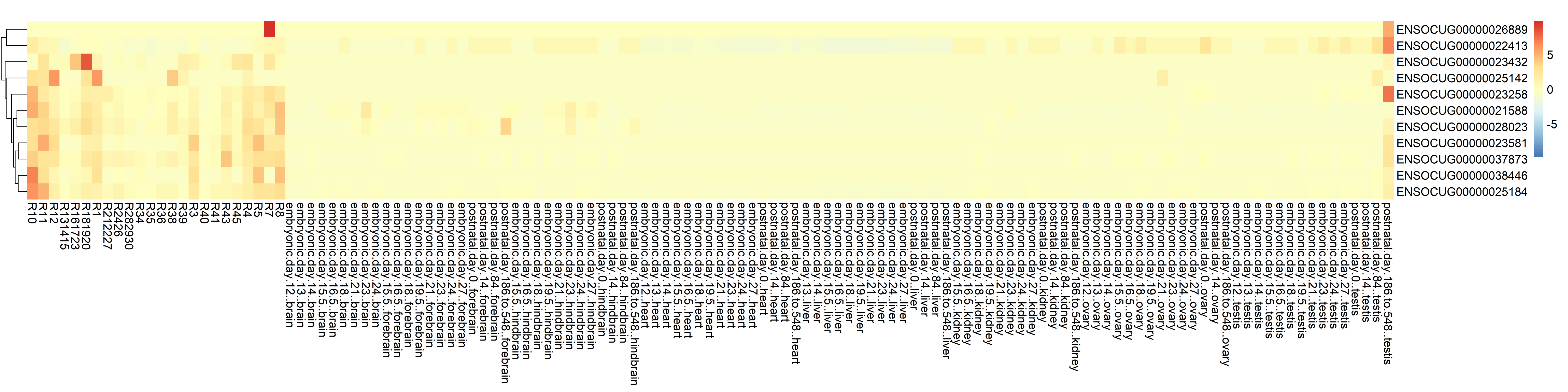
